## Supplementary Figures for "The Ycf48 accessory factor occupies the site of the oxygen-evolving manganese cluster during photosystem II biogenesis"

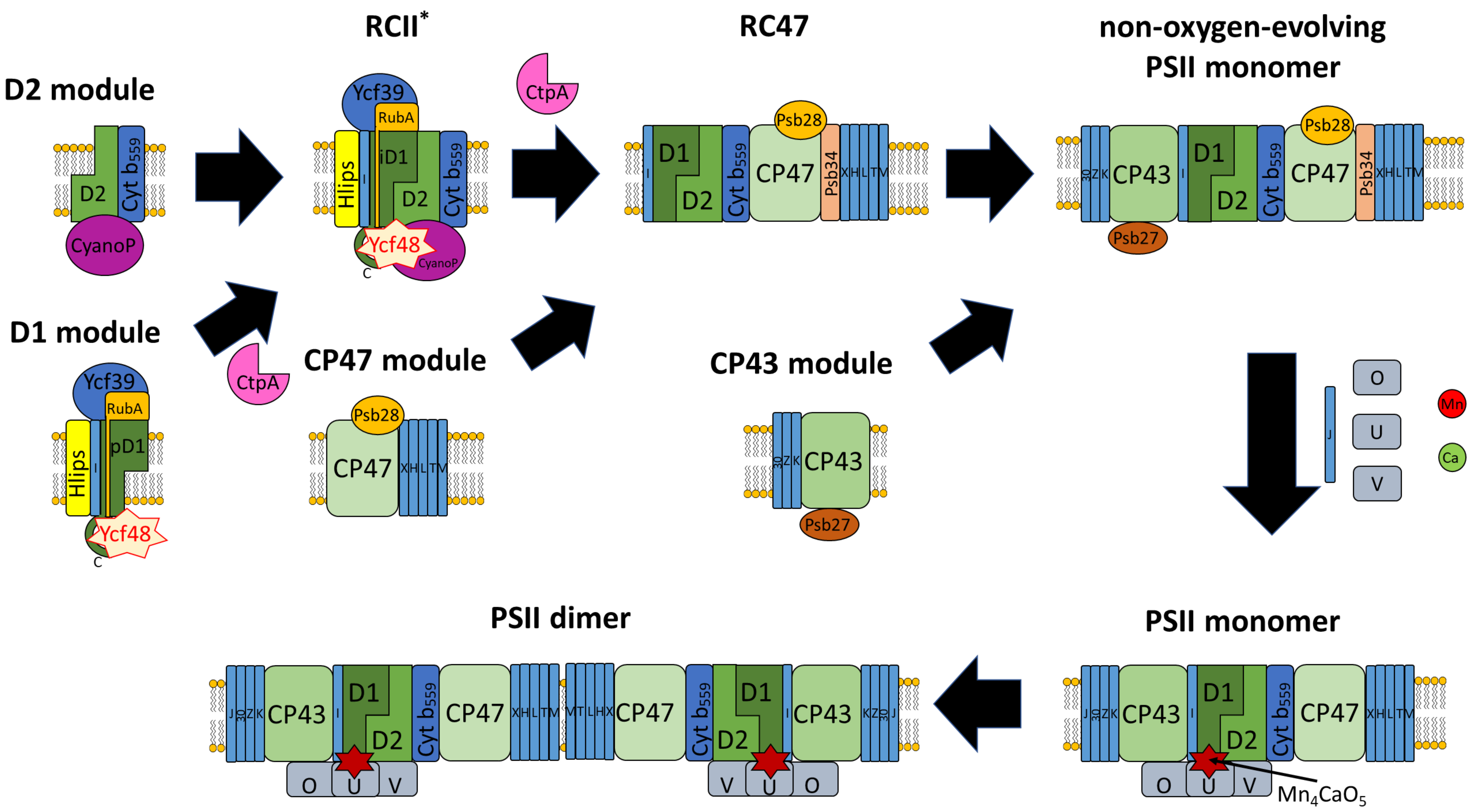


**Supplementary Fig. 1 Model for assembly of PSII in cyanobacteria.** The D1 module (consisting of precursor D1 (pD1), PsbI, Ycf48, RubA and attached Ycf39/Hlips complex) combines with the D2 module (consisting of D2, cytochrome *b*-559 (Cyt $b_{559}$) and CyanoP) to form the PSII reaction center assembly complex, RCII*. Subsequent addition of the CP47 module to form the RC47 complex is followed by addition of the CP43 module to form the non-oxygen-evolving PSII monomer, which can then assemble the $Mn_4CaO_5$ oxygen-evolving cluster and bind PsbJ and the PsbO, PsbU and PsbV extrinsic proteins to form the oxygen-evolving monomeric PSII complex which can then dimerise. PSII subunits are shown without the Psb designation. CtpA is the D1 C-terminal processing protease. Ycf48, CyanoP, RubA, Psb27, Psb28, Psb34 are accessory proteins involved at specific stages of assembly but absent from the final active PSII complex.

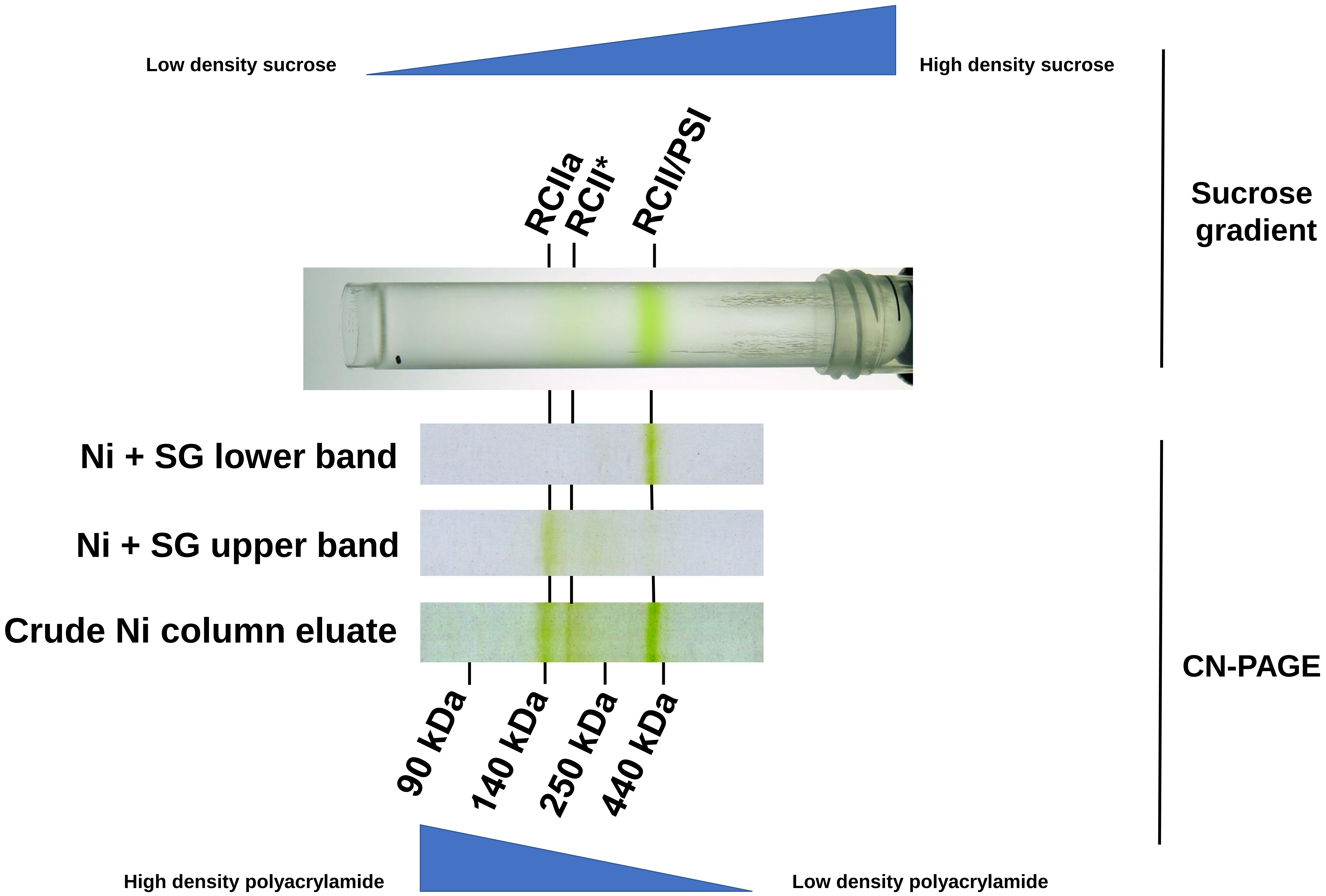


**Supplementary Fig. 2 Isolation of the RCII/PSI complex.** His-tagged D2 complexes isolated by Ni-affinity chromatography from a strain of Syn6803 lacking CP47 (Ni) were subjected to sucrose-density-gradient centrifugation (Sucrose gradient). The upper band in the sucrose gradient contains the (Ni+SG upper band) RCII/PSI complexes analyzed by cryo-EM while the lower band (Ni+SG lower band) is a mixture of RCIIa and RCII*. The purity of the samples was assessed by clear-native PAGE (CN-PAGE) in the presence of n-dodecyl-β-D-maltoside (DM) and sodium deoxycholate (see methods). Approximately 1 µg of Chl of the crude Ni column preparation was loaded onto the sucrose gradient and the CN gel and 0.5 µg of Chl was loaded onto the CN gel for both the lower sucrose gradient band (Ni + SG upper band) containing the RCII/PSI complex and the upper sucrose gradient band (Ni + SG lower band).

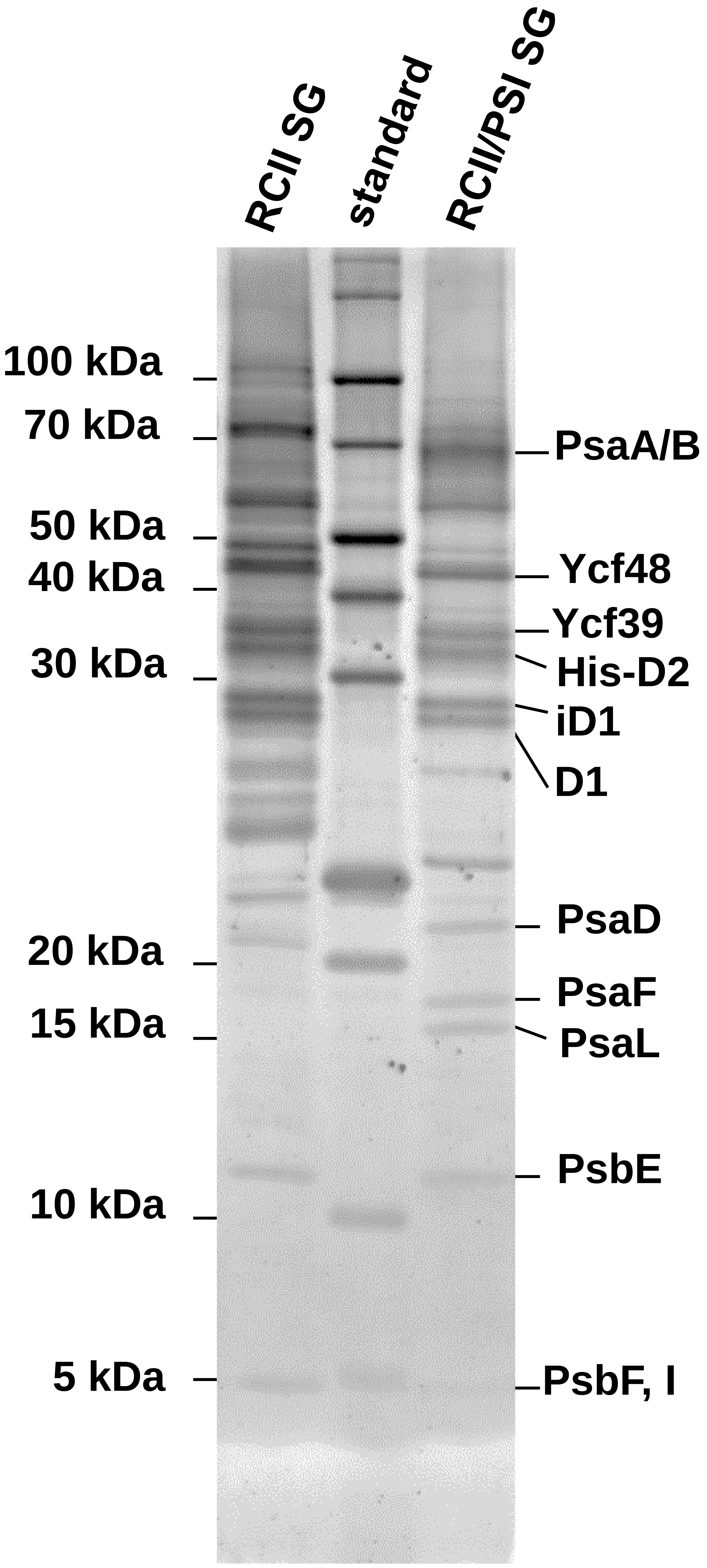


**Supplementary Fig. 3 SDS-PAGE analysis of the upper sucrose gradient fraction containing RCII complexes (RCII SG) and the lower sucrose gradient fraction containing the RCII/PSI complex (RCII/PSI SG).** Proteins identified by mass spectrometry and immunoblotting are indicated. 0.5 µg of Chl was loaded for each lane. Not all proteins detected by mass spectrometry are assigned to a stained band.

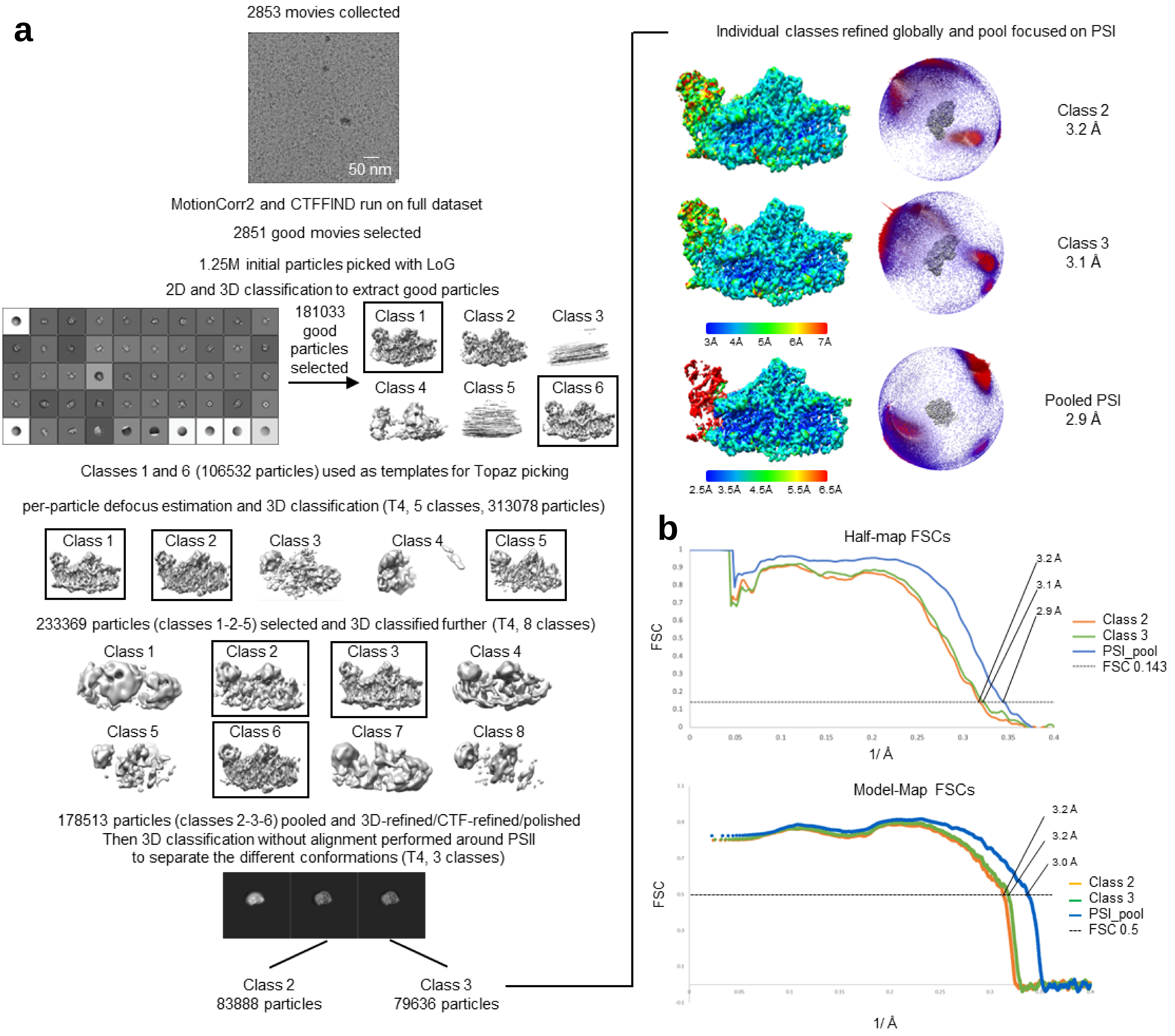


**Supplementary Fig. 4 Processing pipeline, FSC curves and representative densities from the cryo-EM maps**. **a,** Scheme of the processing pipeline, with representative micrograph, pre-processing, classification steps and final maps, coloured by local resolution, with relative angular distribution plot and representative densities. **b,** Gold-standard half-map and model-map Fourier Shell Correlation (FSC) curves for the datasets.

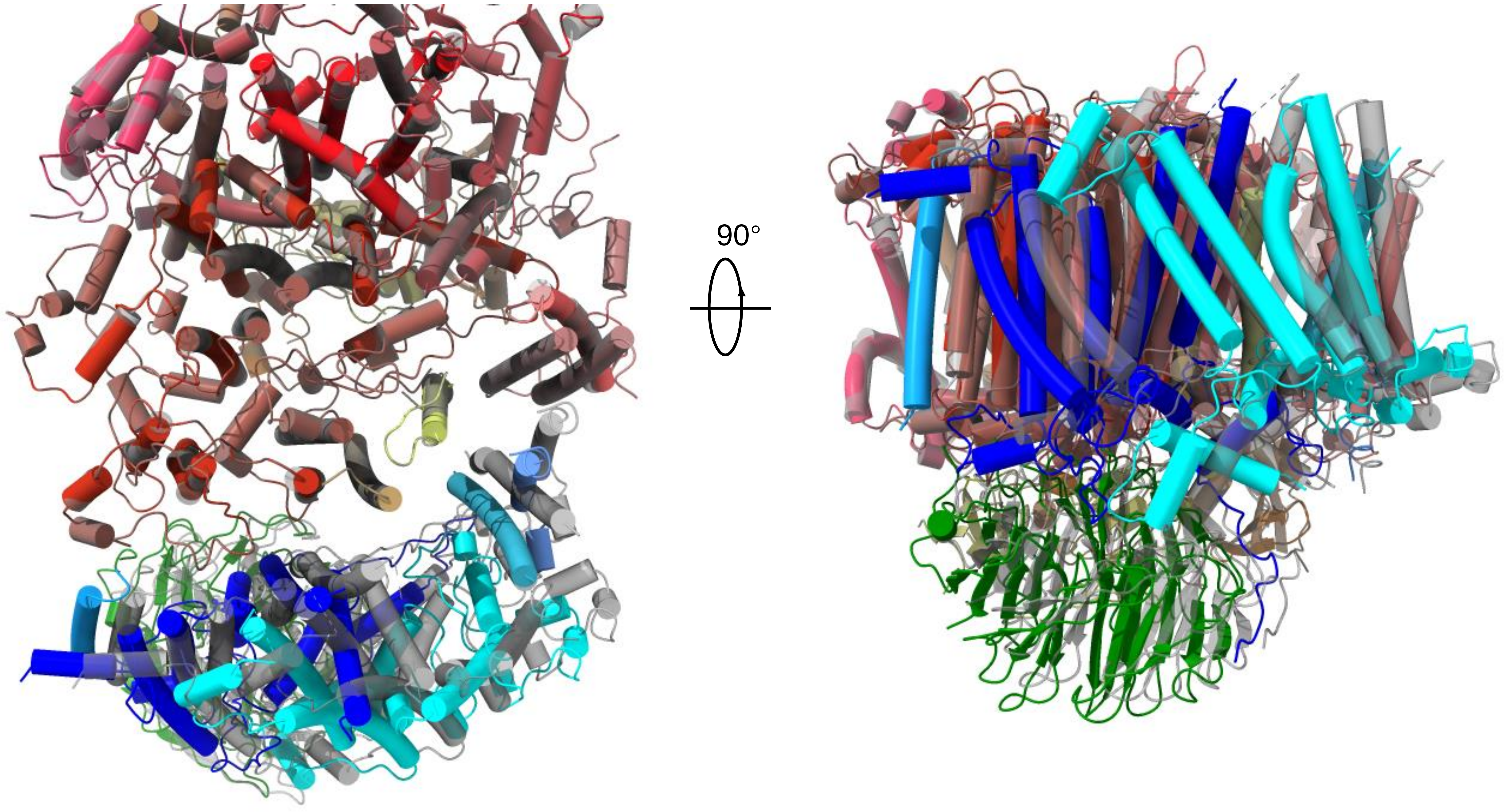


**Supplementary Fig. 5 RCII position shifts in the lower resolution structure.** The lower resolution structure (3.2 Å, shown as grey model) was superimposed to the higher resolution structure used for analysis in the manuscript (shown as coloured model) on PSI: the shift of RCII position, shown from the top view (cytoplasm) on the left and from the side view on the right, is visualized. The positions of D1 (blue), D2 (cyan) and Ycf48 (green) are shown.

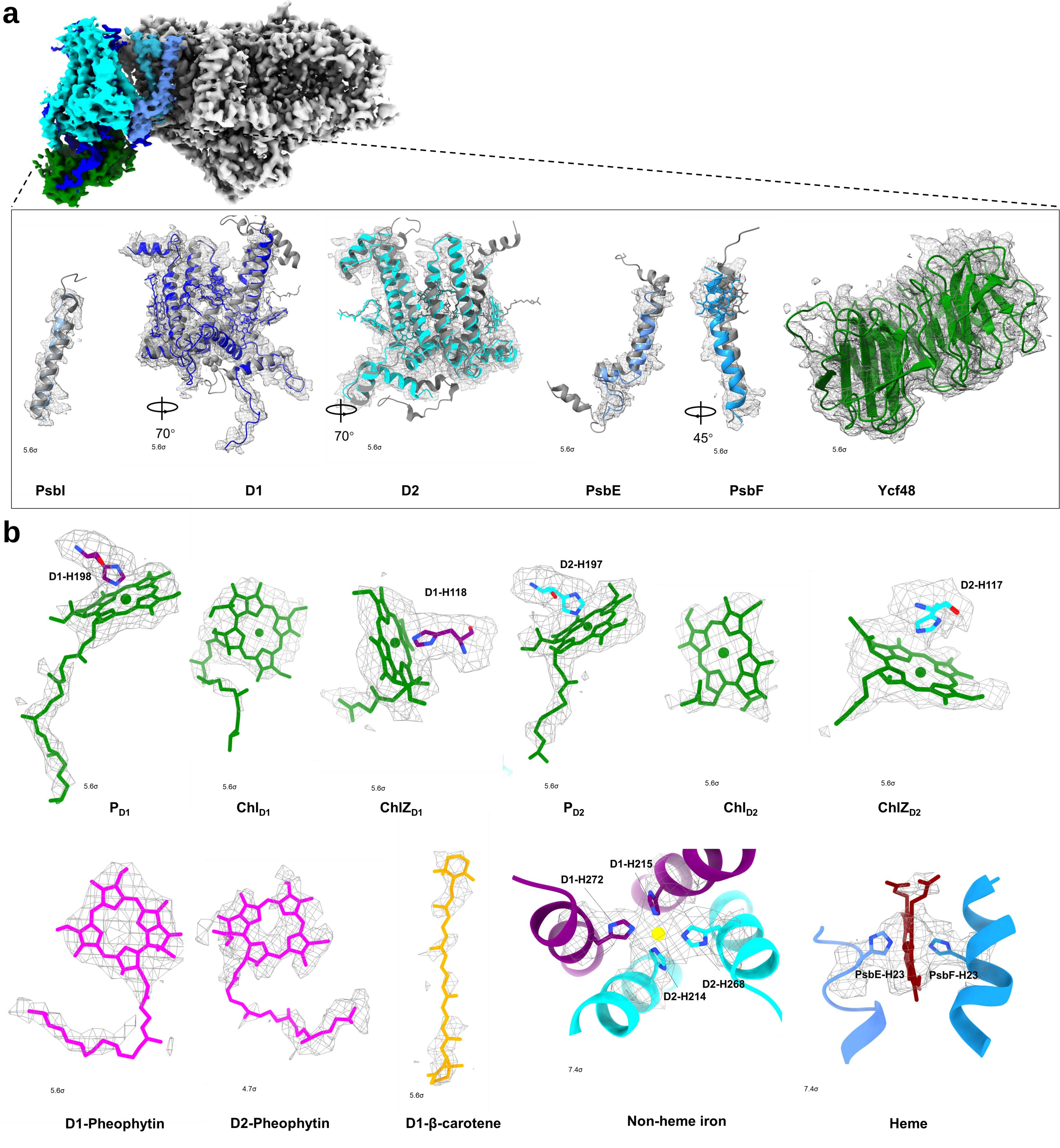


**Supplementary Fig. 6 Class3 Cryo-EM densities and structural models of RCII subunits (a) and various cofactors (b).** Modelled RCII subunits are in colour and in grey the equivalent subunits found in Syn6803 oxygen-evolving PSII (PDB ID: 7N8O. Representative density of subunits are contoured at 5.6σ. Chlorophylls, D1-pheophytin and D1-β-carotene are contoured at 5.6σ. D2-pheophytin is contoured at 4.7σ. Non-heme iron and heme are contoured at 7.4σ.

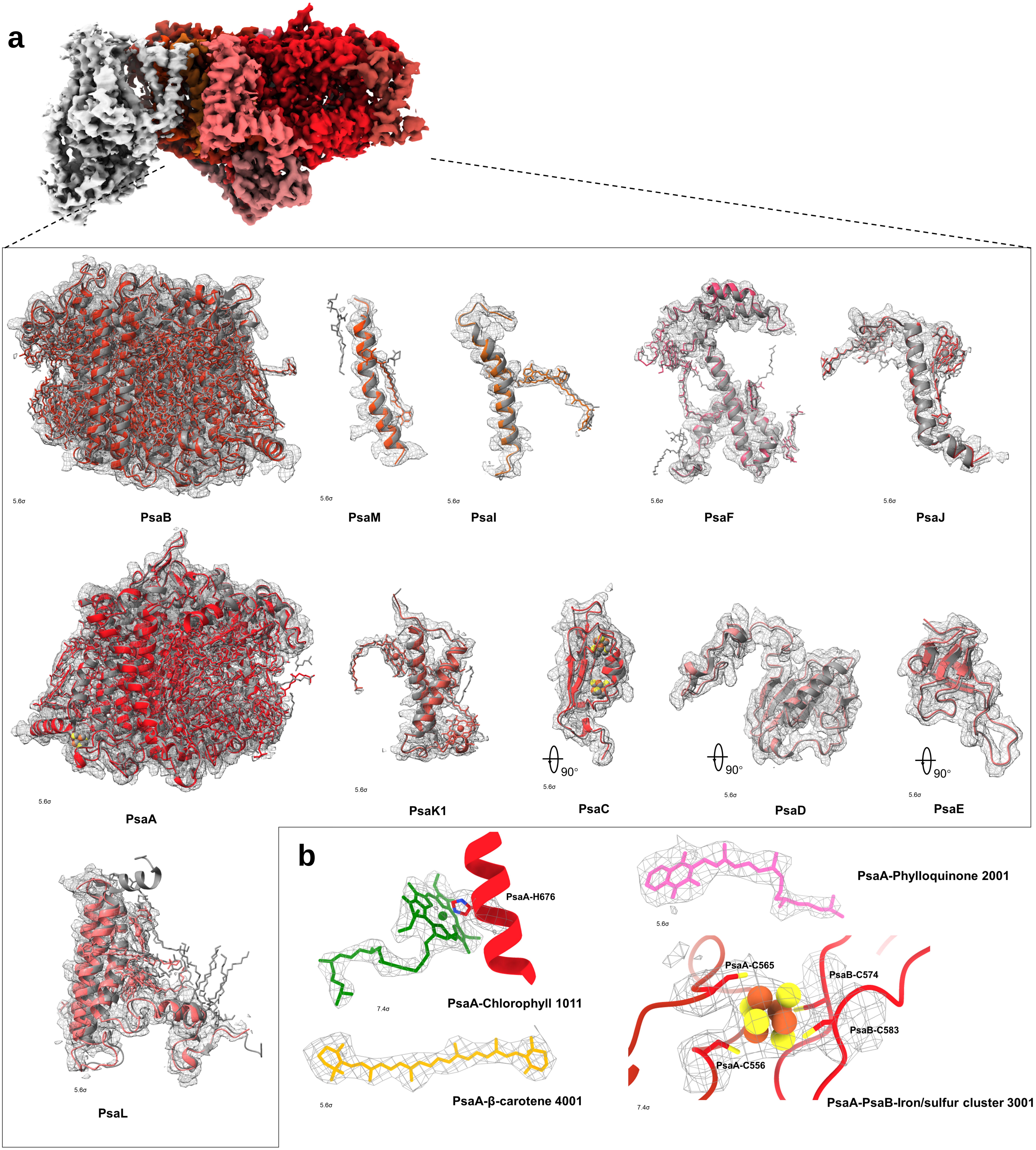


**Supplementary Fig. 7 Class3 Cryo-EM densities and structural models of PSI subunits (a) and selected cofactors (b) in the PSI/RCII complex.** Modelled subunits are shown in colour and in grey the equivalent found in the Syn6803 PSI monomer (PDB ID: 5OY0). Representative density of PSI subunits, PsaA-Chlorophyll 1011, PsaA-Phylloquinone 2001 are contoured at 5.6σ. PsaA-β-carotene 4001 and PsaA-PsaB-Iron/sulfur cluster 3001 are contoured at 7.4σ.

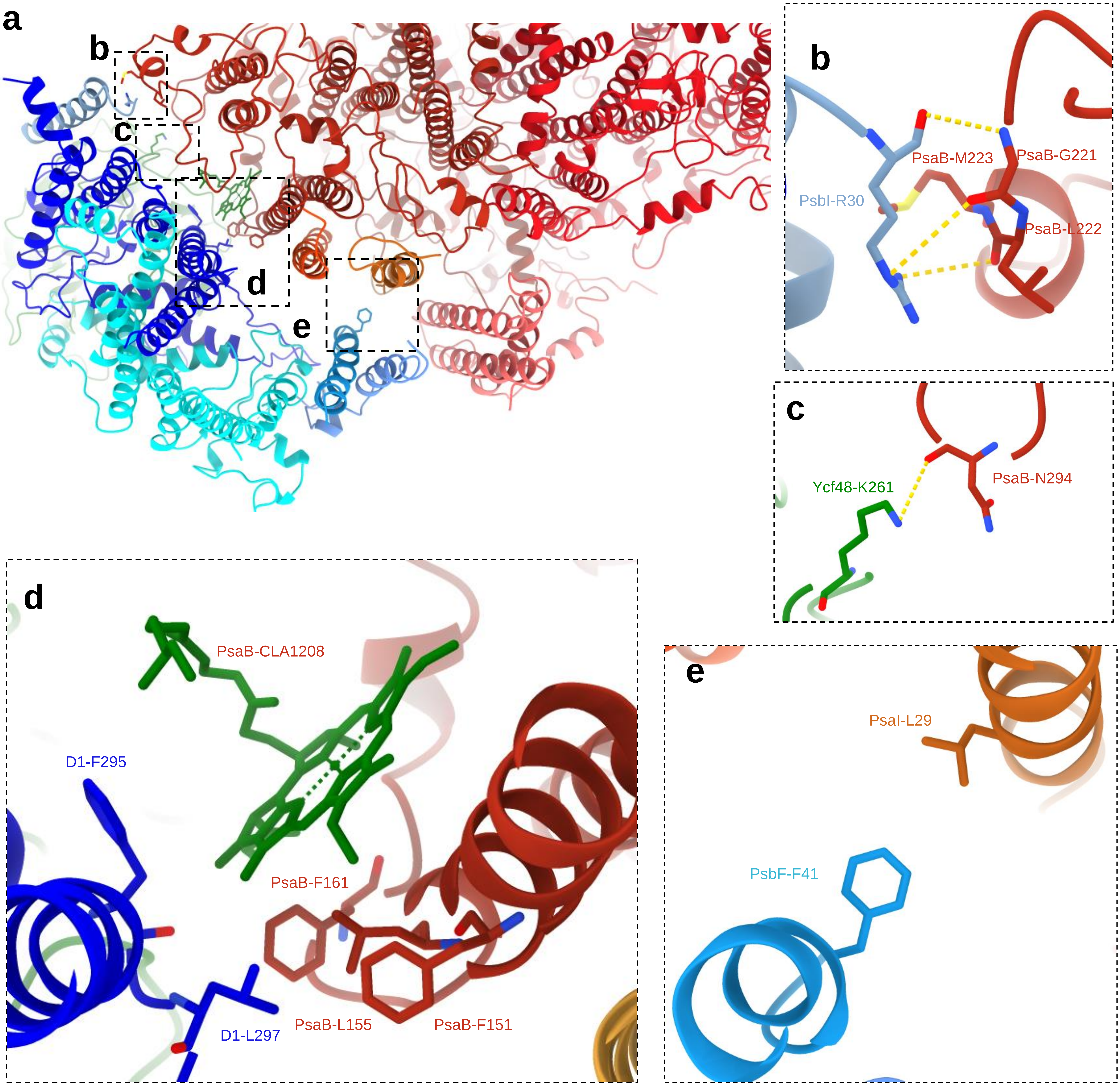


**Supplementary Fig. 8 Interactions between RCII and PSI in the RCII/PSI complex. a,** View of the RCII/PSI complex showing the regions at the interface of RCII and PSI under analysis. **b,** The PsbI-Arg30 backbone carbonyl oxygen, at the N-terminal end of PsbI, potentially H-bonds with the PsaB-Gly221 backbone nitrogen towards the end of the third helix of PsaB. **c,** Ycf48-Lys261, which is at the end of the fifth blade, may H-bond with PsaB-Asn294, a luminal residue at the end of the fourth PsaB helix. **d,** Possible hydrophobic interaction between D1 residues, D1-Phe295 and D1-Leu297, and PsaB residues, PsaB-Phe151, PsaB-Leu155, PsaB-Phe161 and chlorophyll 1208. **e,** C-terminal PsbF residue, PsbF-Phe41, may have hydrophobic interaction with PsaI-Leu29.

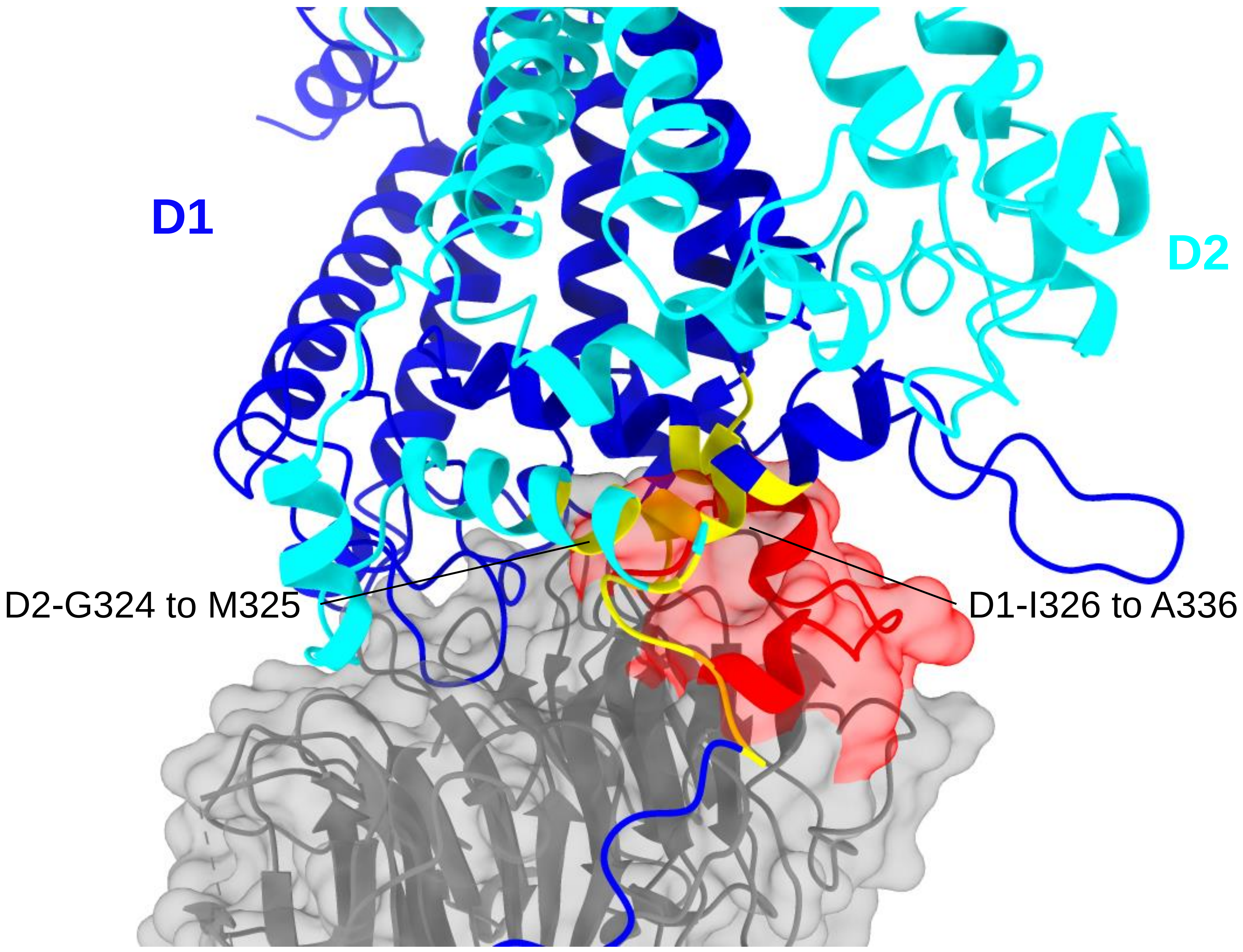


**Supplementary Fig. 9 The Ycf48 eukaryotic loop clashes when modelled into the cyanobacterial RCII complex.** Structure of *Cyanidioschyzon merolae* Ycf48 (PDB ID: 5OJ3, shown in grey) was superimposed onto Syn6803 Ycf48 in the PSI/RCII complex. The 19 amino-acid-residue insertion in the eukaryotic loop is coloured red. D1 is shown in blue and D2 in cyan. Sites of steric clash (D1-Ile326 to D1-Ala336 and D2-Gly324 to D2-Met325) are coloured yellow.

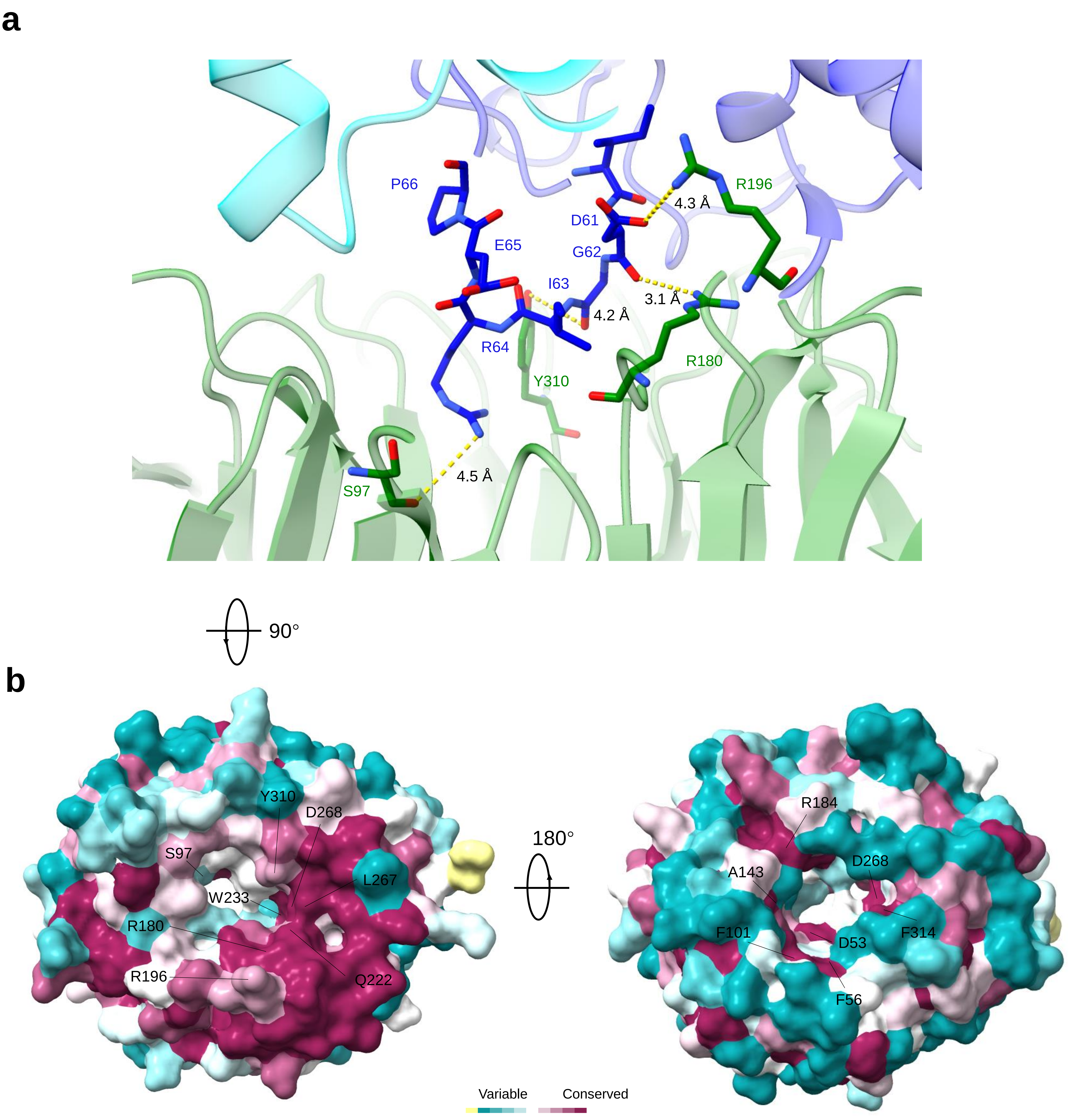


**Supplementary Fig. 10 The Ycf48 channel. a,** D1 loop at the exit of the Ycf48 channel. D1 in blue, Ycf48 in green and D2 in cyan. b, Identification of conserved residues within the Ycf48 channel viewed from the exit (left) and the entrance (right). Purple shows the most conserved regions and blue shows more variable regions. There is insufficient data to show the Ycf48-His31, which is not conserved based on IJMS analysis (ref), conservation score by ConSurf and is shown in yellow.

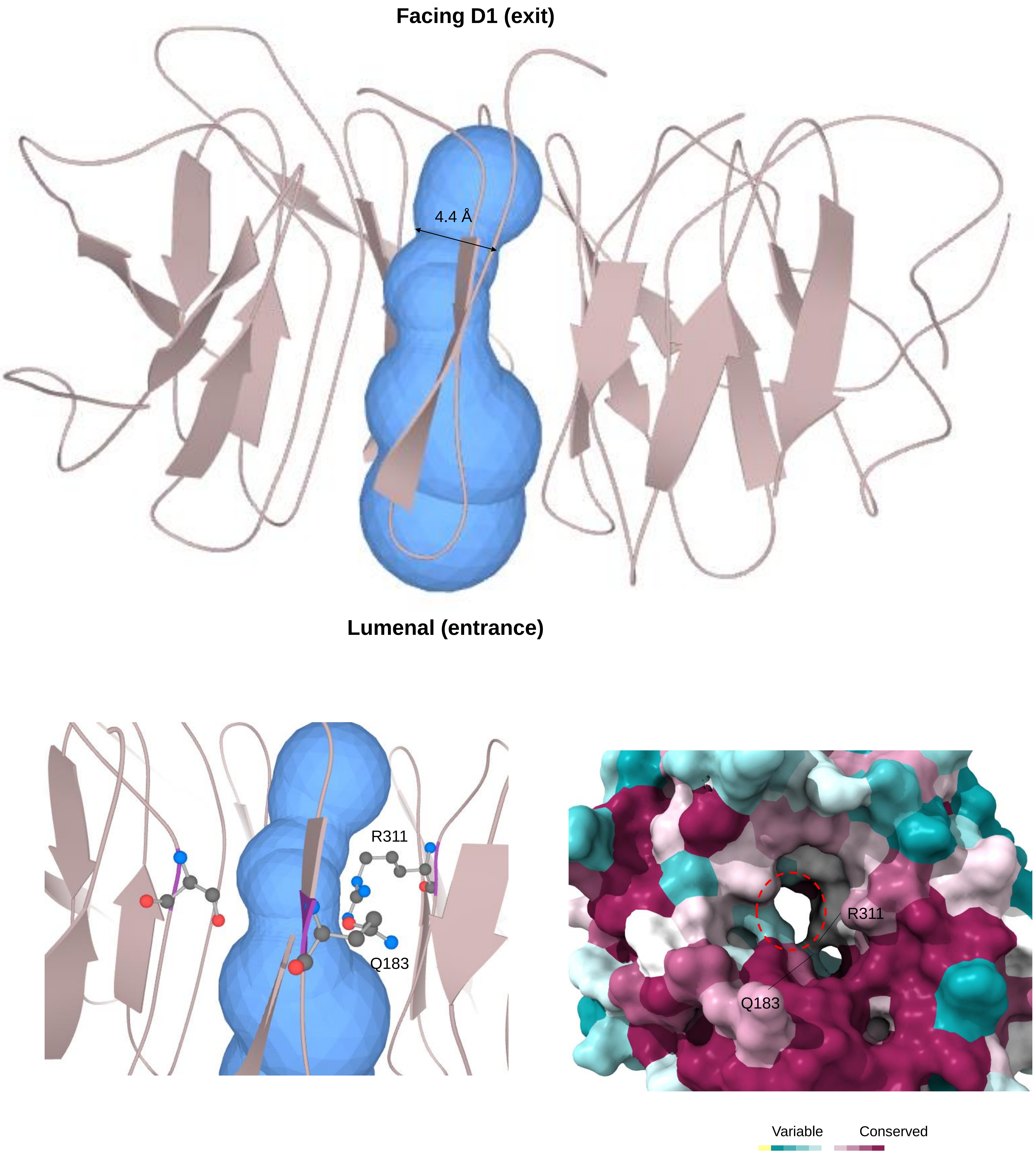


**Supplementary Fig. 11 Ycf48 central channels.** Ycf48 channels were analysed by MOLEonline (https://mole.upol.cz/). The central hole is split by R311 and Q183 into two tunnels, the bottleneck diameter of the widest tunnel is shown. Lumenal side of the channel is defined as the ‘entrance’ and the side facing D1 is the ‘exit’. Minor tunnels not shown.

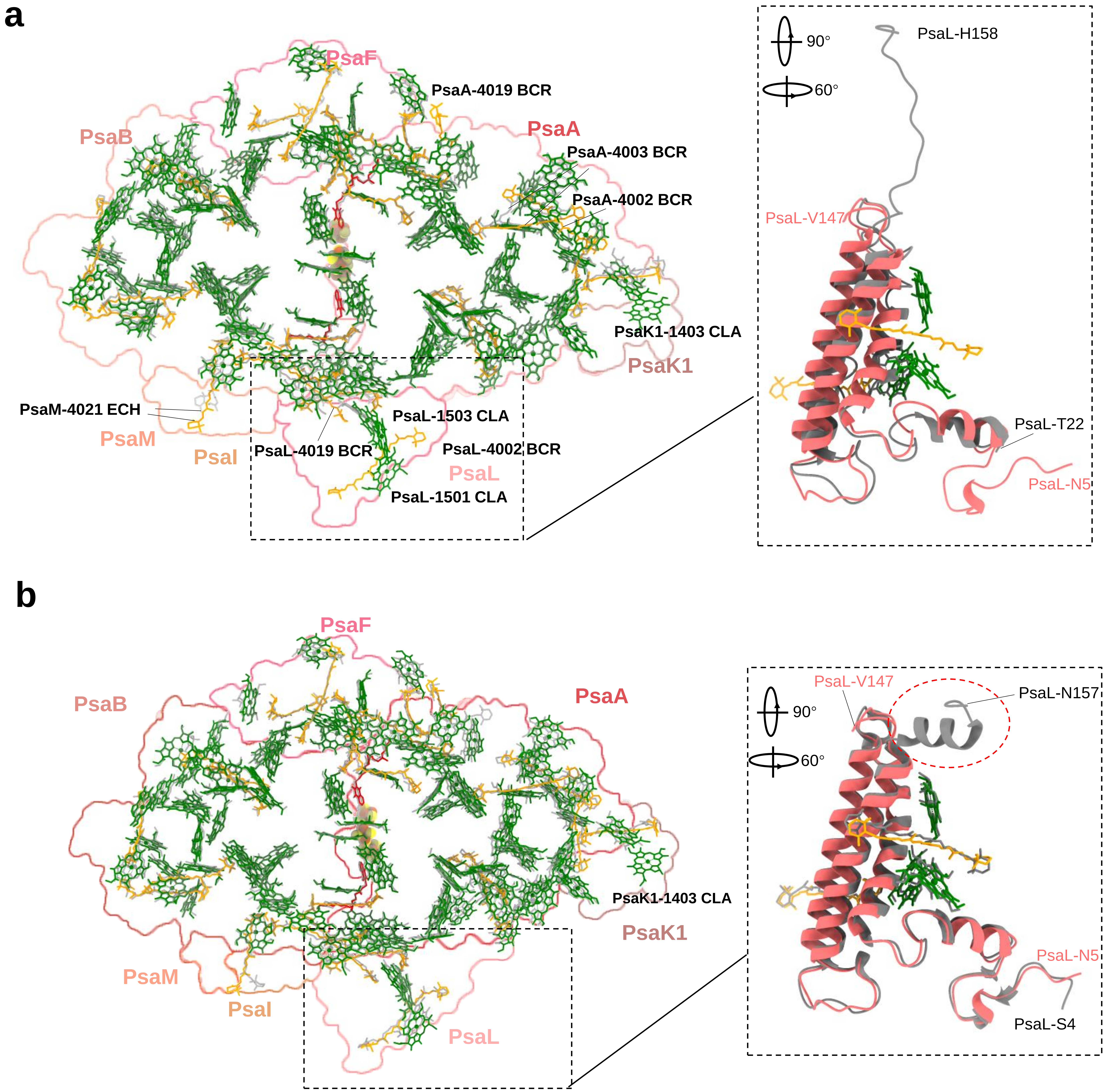


**Supplementary Fig. 12 Comparison of pigments PSI in the RCII/PSI complex with published structures of monomeric and trimeric PSI from Syn6803. a,** Top view comparison of PSI cofactors in the PSI/RCII complex and the Syn6803 PSI monomer (PDB ID: 6HQB). Chlorophylls are shown in green, carotenoids in yellow and phylloquinone in red. Three additional chlorophylls (CLA) and four additional β-carotenes (BCR) are indicated in bold. One PsaM ehinenone (PsaM-4021 ECH) and one PsaA β-carotenes (PsaA-4003 BCR) show different positions. The dashed box shows differences between PsaL in the PSI/RCII complex (pink) and PSI monomer (grey). **b,** Top view comparison of PSI cofactors in the RCII/PSI complex and the monomer found in the Syn6803 PSI trimer (PDB ID: 5OY0). Dashed box compares the structures of PsaL in the RCII/PSI complex (coloured pink) and in the trimer (in grey). The short C-terminal helix of PsaL, which is involved in PSI trimerization, cannot be modelled in the PSI/RCII structure. Additional chlorophyll indicated in bold.

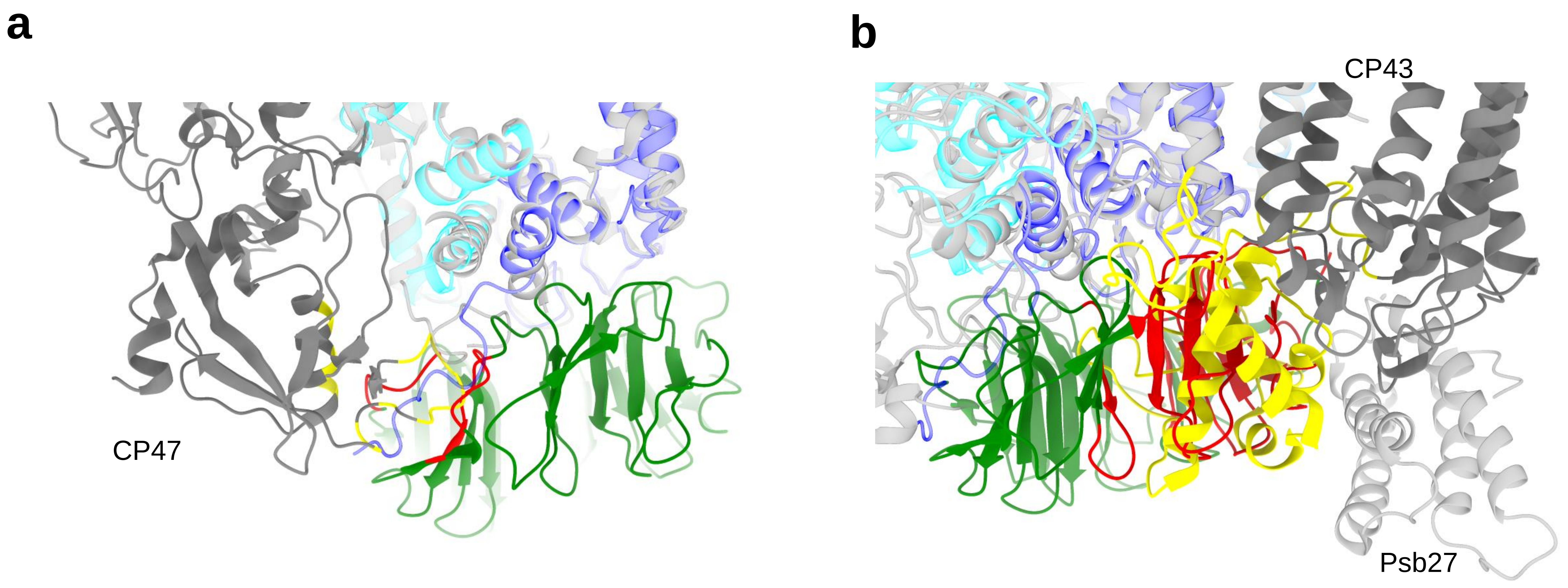


**Supplementary Fig. 13 Potential steric clashes between CP47 and CP43 when the structure of Ycf48 is inserted into the structures of RC47 and the Psb27/PSII complex. a,** Loop region of CP47 (CP47-Pro382 to CP47-Lys389, in yellow) and several luminal helix residues (CP47-Ser391, Glu393, Gln394, Pro415, Lys418 and Arg422, in yellow) clash with blade 3 residues of Ycf48 (Ser131-Ala137, Asp154-Tyr159, Ala170, Glu173-Ala175, in red) when *T. vulcanus* RC47 (PDB ID:7DXA) is superimposed on the structure of RCII. Ycf48 is shown in green. D1 and D2 in the RCII/PSI complex are shown in transparent blue and cyan, respectively. RC47 subunits are shown in grey, CP47 is shown in solid and other subunits are transparent. **b,** *T. vulcanus* Psb27/PSII (PDB ID:7CZL) was superimposed onto RCII. Psb27/PSII subunits are shown in grey, CP43 is solid and other subunits are transparent. Ycf48 is shown in green. D1 and D2 in RCII/PSI complex are shown in transparent blue and cyan, respectively. The large lumenal region of CP43 (Pro217-Asn228, Pro304-Val420, in yellow) clashes with blades 4, 5 and 6 of Ycf48 (His31, Arg180-Arg189,Tyr200, Asn214-Asp299, Tyr310-Phe314, in red).

| | Subunit name and Uniprot code | Size (Da) Length (AA) | MS Intensity | Blot | Chain name in the model | Range built/total residues | Unmodeled residues | % modelled residues | Cofactors | Notes |
|---|---|---|---|---|---|---|---|---|---|---|
| RCII | D1, PsbA P16033 | 39695 360 | 90237000 | + | A | 13-224,267-344/1-344 | 1-12, 225-266 | 84.3 | chlorophyll, non-heme iron, β-carotene, pheophytin | Cytoplasmic loop around non-heme iron unmodeled |
| | D2, PsbD P09192 | 39466 352 | 112990000 | + | D | 32--222,244-330/1-352 | 1-31, 223-243,331-352 | 79.0 | chlorophyll, non-heme iron, pheophytin | Cytoplasmic loop around non-heme iron unmodeled |
| | PsbE P09190 | 9442 81 | 28092000 | + | E | 20-56/2-81 | 2-19, 57-81 | 46.3 | heme | N-terminal and C-terminal short helices unmodeled |
| | PsbF P09191 | 4929 44 | 26176000 | + | F | 17-44/2-44 | 2-16 | 65.1 | heme | N-terminal tail unmodeled |
| | PsbI Q54697 | 4306 38 | n.d. | + | I | 2-30/1-38 | 1, 31-38 | 76.3 | | C-terminal tail unmodeled |
| | Ycf48 P73069 | 37267 342 | 279630000 | + | S | 31-333/29-339 | 29-30, 334-339 | 97.1 | | Ycf48 is lipidated at Cys29, three C-terminal residues are removed |
| | RubA P73068 | 12570 115 | 31088000 | ± | | | | | | |
| | Ycf39 P74429 | 36496 326 | 11390000 | ± | | | | | | |
| | HliD P73563 | 6472 57 | 630300 | ± | | | | | | |
| | RCII | | | | | 965/1168 | | 82.6 | | |
| PSI | PsaA P29254 | 82950 751 | 26052000 | + | a | 11-751/1-751 | 1-10 | 98.7 | chlorophyll, phylloquinone, β-carotene, $Fe_4S_4$ cluster | |
| | PsaB P29255 | 81292 731 | 8619200 | + | b | 3-731/2-731 | 2 | 99.9 | chlorophyll, phylloquinone, β-carotene, $Fe_4S_4$ cluster, canthaxanthin | |
| | PsaC P32422 | 8828 81 | 9414800 | ± | c | 2-81/2-81 | | 100.0 | $Fe_4S_4$ cluster | |
| | PsaD P19569 | 15644 141 | 55535000 | + | d | 3-141/2-141 | 2 | 99.3 | | |
| | PsaE P12975 | 8145 74 | 25510000 | - | e | 2-70/2-74 | 71-74 | 94.5 | | |
| | PsaF P29256 | 18249 165 | 58829000 | + | f | 24-165/24-165 | | 100.0 | chlorophyll, zeaxanthin | 1-23 is signal peptide |
| | PsaI | | | | i | 1-40/1-40 | | 100.0 | β-carotene, 3 'OH echinenone | |
| | PsaJ | | | | j | 1-40/1-40 | | 100.0 | chlorophyll, β-carotene, zeaxanthin | |
| | PsaK1 P72712 | 8644 86 | 335670 | n.a. | k | 9-83/1-86 | 1-8, 84-86 | 87.2 | chlorophyll, β-carotene | |
| | PsaK2 P74564 | 9306 90 | 2946400 | n.a. | | | | | | |
| | PsaL P37277 | 16624 157 | 27715000 | + | l | 5-147/1-157 | 1-4, 148-157 | 91.1 | chlorophyll, β-carotene | C-terminal helix unmodeled |
| | PsaM P72986 | 3380 31 | 3305200 | n.a. | m | 1-31/1-31 | | 100.0 | echinenone | |
| | PSI | | | | | 2229/2270 | | 98.2 | | |
| Total | RCII/PSI | | | | | 3194/3438 | | 93.0 | | |

**Supplementary Table 1 List of PSII and PSI subunits detected in the RCII/PSI complex by mass spectrometry (MS) and immunoblotting (Blot) and subunit composition of the deposited model.** The MS analysis of proteins precipitated from the preparation was performed using NanoElute UHPLC (Bruker) on-line coupled to a high-resolution mass spectrometer (Bruker Impact HD). n.d. means not detected, n.a. means not analysed.

| | #1 class2 (EMDB-X (PDB X) | #1 class3 (EMDB-X) (PDB X) | #3 PSI_focus_ref (EMDB-X) (PDB X) |
|---|---|---|---|
| **Microscope** | Glacios | Glacios | Glacios |
| **Camera** | Falcon 3 linear | Falcon 3 linear | Falcon 3 linear |
| **Magnification** | 120000X | 120000X | 120000X |
| **Voltage (kV)** | 200 | 200 | 200 |
| **Electron exposure (e–/Å$^2$)** | 91 | 91 | 91 |
| **Automation software** | EPU | EPU | EPU |
| **Number of frames** | 58 | 58 | 58 |
| **Defocus range (μm)** | c.a. -1.2 to -2.5 | c.a. -1.2 to -2.5 | c.a. -1.2 to -2.5 |
| **Pixel size (Å)** | 1.22 | 1.22 | 1.22 |
| **Symmetry imposed** | C1 | C1 | C1 |
| **Number of micrographs** | 2853 | 2853 | 2853 |
| **Initial particle images (no.) from Topaz** | 313078 | 313078 | 313078 |
| **Final particle images (no.)** | 83888 | 79636 | 163524 |
| **Map resolution (Å) at 0.143 FSC threshold** | 3.2 | 3.1 | 2.9 |
| **Local resolution range (Å)** | 2.7-9.5 | 2.7-9.2 | 2.7-5.0 |
| **Refinement** | | | |
| **Initial model used (PDB code)** | RCII (6WJ6); PSI (5OY0); Ycf48 (2XBG) | RCII (6WJ6); PSI (5OY0); Ycf48 (2XBG) | (5OY0) |
| **Refinement package** | Phenix, real space | Phenix, real space | Phenix, real space |
| **Model resolution (Å) at 0.5 FSC threshold** | 3.2 | 3.2 | 3.0 |
| **Cross-correlation** | | | |
| **Mask** | 0.86 | 0.87 | 0.89 |
| **Volume** | 0.85 | 0.86 | 0.88 |
| **Map sharpening B factor (Å$^2$)** | -60.2 | -55.3 | -54.6 |
| **Model composition** | | | |
| **Non-hydrogen atoms** | 32740 | 32747 | 24597 |
| **Protein residues** | 3198 | 3194 | 2230 |
| **Ligands** | 145 | 146 | 134 |
| **B factors (Å$^2$) (mean)** | | | |
| **Protein** | 89.15 | 93.00 | 69.14 |
| **Ligand** | 66.23 | 69.11 | 65.03 |
| **R.m.s deviations** | | | |
| **Bond lengths (Å)** | 0.003 | 0.003 | 0.002 |
| **Bond angles (°)** | 0.583 | 0.573 | 0.597 |
| **Validation** | | | |
| **MolProbity score** | 1.86 | 1.82 | 1.87 |
| **EMRinger score** | 2.66 | 2.62 | 2.58 |
| **Clashscore** | 14.79 | 13.81 | 13.55 |
| **Poor rotamer (%)** | 0.08 | 0.08 | 1.34 |
| **C-beta outliers %** | 0 | 0 | 0 |
| **CaBLAM outliers (%)** | 1.54 | 1.54 | 1.51 |
| **Ramachandran plot** | | | |
| **Favored (%)** | 96.93 | 96.99 | 97.28 |
| **Allowed** | 3.07 | 3.01 | 2.72 |
| **Disallowed (%)** | 0 | 0 | 0 |

**Supplementary Table 2 Cryo-EM data collection, refinement and validation statistics**
